# Dicarboxylic acids synergize with yeast and human Hsp60/10 systems to mimic GroEL/ES

**DOI:** 10.1101/2024.05.15.594092

**Authors:** Deepika Gautam, Mohammad Aaquib, Manisha Kochar, Kausik Chakraborty

## Abstract

GroEL/ES has been the archetype to understand the function of the class I chaperonins (Hsp60/10 systems). While very similar in structure, the human or yeast mitochondrial one has diminished negative charge density in the central cavity. These chaperones had also lost their ability to assist a substrate of *E.coli* GroEL/ES. Here, we show that the eukaryotic Hsp60/10 systems can synergize with dicarboxylic acids in vitro at the physiological concentration of these metabolites to mimic the activity of *E. coli* GroEL/ES. Combining these Hsp60/10s and metabolites effectively alters the folding landscape like GroEL/ES; this is specific for the eukaryotic chaperonins and not the prokaryotic homologs with less negatively charged cavities. Thus, we identify a potential cooperation between molecular and chemical chaperones that may have important physiological implications linking metabolism to proteostasis.

---

Proteins are functional in their native states. While most can reach their native state without assistance, many require additional help. Multiple proteins and small molecules are known to aid in assisting proteins to reach their native state; these are known as molecular chaperones and chemical chaperones, respectively [1],[2],[3],[4],[5],[6]. One of the most conserved molecular chaperones is the class I chaperonin, Heat Shock Protein 60 (Hsp60). It acts with its co-chaperone Heat Shock Protein 10 (Hsp10) to form an ATP-dependent folding machinery (Hsp60/10 chaperone system) [7]. Heptamers of Hsp60 form a barrel; two such barrels are arranged back-to-back to generate the canonical tetradecameric assembly of Hsp60. Hsp10s form a heptameric lid that closes the cavity of Hsp60 in an ATP-hydrolysis-dependent manner [8],[9],[10],[11].

*E. coli* derived GroEL/ES is the most studied Hsp60/10 system; it has been used extensively to understand the role of Hsp60/10 systems in vivo [12],[13],[14],[15],[16],[17]. While this has led to extensive insights into the mechanism(s) through which this chaperone assists in folding its substrates, recent literature suggests that there may be ortholog-specific mechanisms that differ from the GroEL/ES system. Orthologs differ in negative charge density inside the Hsp60/10’s barrel – this leads to a divergence in the ability of the chaperonins to alter the folding landscape of its client [18]. While the difference is not unexpected, given that these systems adapt to their proteome, little is known about their specializations.

The eukaryotic class I chaperonins are found in the mitochondria and the chloroplast. The mitochondrial Hsp60/10s from most model eukaryotes have a less negatively charged central cavity than GroEL/ES [18]. While the proteome of mitochondria is different, the metabolite pool is also significantly different from that of E. coli cytosol [19]. Decades earlier, it was shown that chaperones involved in disaggregating protein aggregates could cooperate with dicarboxylic acids to aid in folding substrate proteins [20]. However, nothing is known about this cooperation with other chaperones or if this cooperation is a conserved phenomenon. Given that it was possible to complement the loss of negative charges in the eukaryotic Hsp60/10 systems by adding negatively charged small molecules, we asked if the eukaryotic chaperonins have evolved mechanisms to cooperate with dicarboxylic acids that are bonafide metabolites found in eukaryotic mitochondria.

We show that eukaryotic Hsp60/10 systems, not the prokaryotic ones, evolved to cooperate with some of the dicarboxylic acids at the reported physiological concentration of these metabolites. While these metabolites allow the chaperonins to aid in folding of the substrates, they do not change the allosteric cycle, suggesting that this cooperation will likely be a complementation effect. We also show that these small molecules synergize with the Hsp60/10 systems to alter the folding landscape like GroEL/ES. This suggests that the yeast and human Hsp60/10s likely function along with the metabolites in vivo and, hence, could have lost their negative charge due to relaxed selection.

## Results and Discussion

### sGFP is a substrate of GroEL/ES in vivo and in vitro but not of the eukaryotic homologs

We recently reported a new model substrate of GroEL/ES derived from Green Fluorescent Protein (GFP)[21]. The mutation that conferred GroEL/ES dependence was a K45E substitution (the mutant is hereafter referred to as sGFP). As measured by the fluorescence of the protein, the activity of this model substrate is easily measurable in vivo. We inserted mCherry as an independent translating unit in the same mRNA encoding GFP using a bicistronic construct; this allowed us to measure the activity of the protein after normalizing for intercellular differences in plasmid copy number, induction levels and other transcriptional and translational differences between cells (Figure 1A). The activity of sGFP increases *in vivo* (in all cases *in-vivo* means in *Escherichia coli* unless otherwise stated) when GroEL/ES is co-expressed (Figure 1B). The refolding rate of the purified protein *in vitro* also increased significantly and dramatically when assisted by the GroEL/ES system (Figure 1C). More importantly, as reported earlier, the rates of refolding of sGFP did not increase in the presence of either *Saccharomyces cerevisiae* Hsp60/10 (yHsp60/10) or human Hsp60/10 (hHsp60/10) (Figure 1C); these lack the heavily negatively charged central cavity found in GroEL/ES [18]. This was recapitulated *in vivo*; co-expression of these chaperonin homologs could not aid the folding of sGFP *in vivo* (Figure 1B), as reported earlier [18].

**Figure 1.**
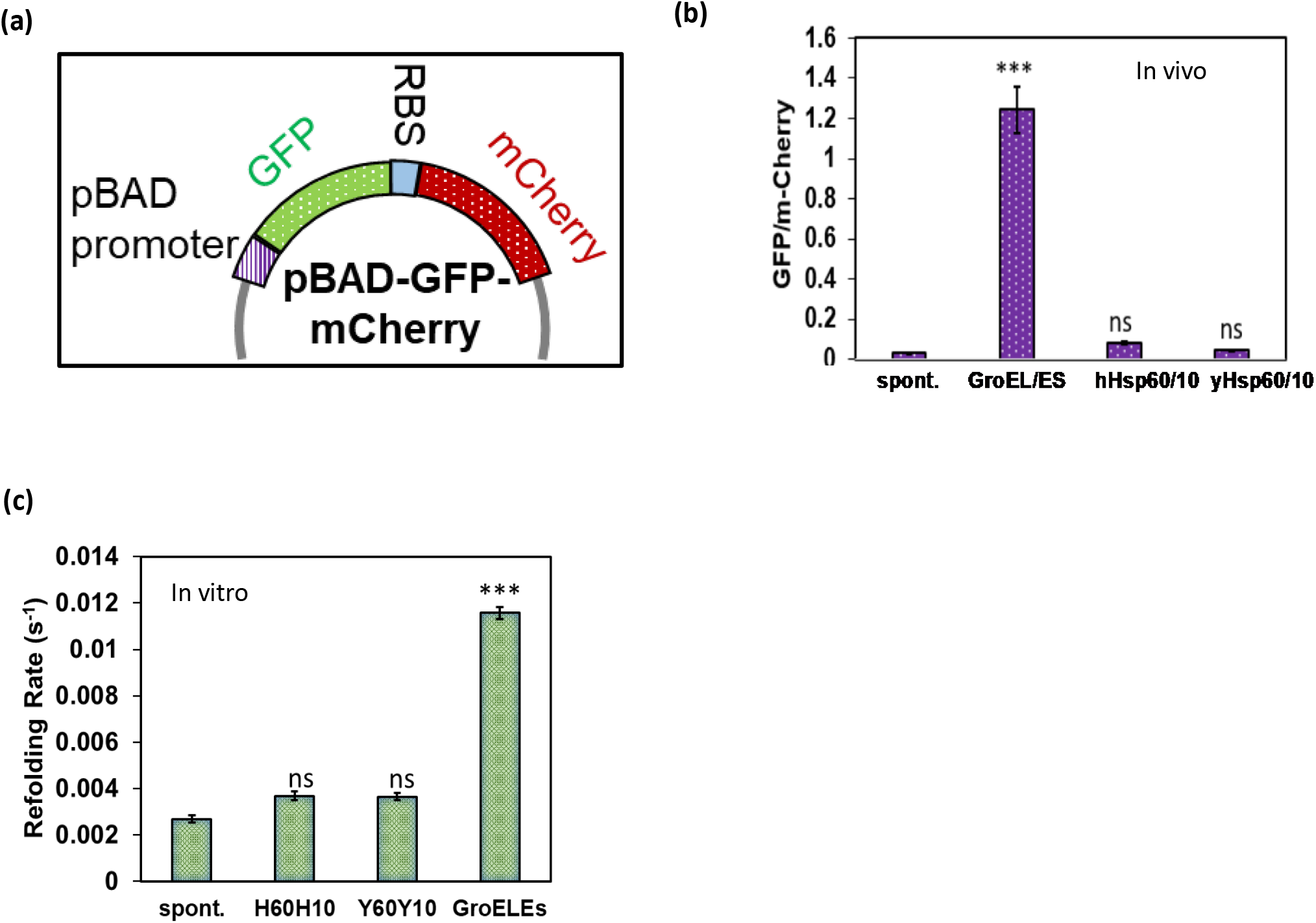
sGFP is a substrate for GroEL/ES both *in-vivo* and *in-vitro*, yet eukaryotic homologs do not recognize it. **(a)** The design bicistronic construct plasmid involves a specific reporter configuration where both Green Fluorescent Protein (GFP) and mCherry are positioned under the control of the same arabinose-inducible pBAD promoter. These two genes are arranged in an operon structure, ensuring they are co-regulated and transcribed. **(b)** Bar plot for *in-vivo* GFP/mCherry fluorescence of sGFP in the presence of overexpression of GroEL/ES, human Hsp60/10, yeast Hsp60/10. **(c)** Comparison of the refolding kinetics of sGFP spontaneous and GroEL/ES Homologs like hHsp60/10, yHsp60/10.

### Dicarboxylic acids potentiate the acceleration with eukaryotic but not prokaryotic chaperonins

Earlier, we had observed that high concentrations of negatively charged small molecules, namely dicarboxylic acids, could accelerate the refolding of sGFP in vitro [18]. Notably, at very high like 100mM concentrations, these small molecules could also assist the rate of hHsp60/10 of yHsp60/10 assisted refolding of sGFP [18]. This indicated that at these high concentrations, negative charges added from outside could partially compensate for the lack of negative charge on the surface of these eukaryotic chaperonins. To investigate the role of these small molecules and the importance of the negatively charged cavity more quantitatively, we focused on four dicarboxylic acids: succinate, fumarate, aspartate, and malate (Figure 2A). At concentrations till 10mM, these small molecules had minimal effect on the *in vitro* refolding rate of sGFP (Figure 2B). Remarkably, these small molecules exhibited high synergism with yHsp60/10 or hHsp60/10 systems at concentrations as low as 0.1mM (Figure 2C-J). While succinate, aspartate, and malate could accelerate yHsp60/10 dependent refolding of sGFP dramatically, fumarate accelerated hHsp60/10 assisted refolding of sGFP. The refolding rate did not monotonically increase with concentration and, in some cases, decreased at high concentrations of the small molecules (Figure-2g, Figure-2h). Notably, neither the chaperonins alone nor the small molecules (as shown by the reference lines in the respective plots) could assist in refolding sGFP at these concentrations. While hHsp60/10 does not assist the refolding of sGFP to any significant extent, in the presence of 0.1mM Fumarate, the refolding rate of sGFP (∼0.09s^-1^) is comparable to GroEL/ES assisted folding of sGFP (∼0.1s^-1^).

**Figure 2.**
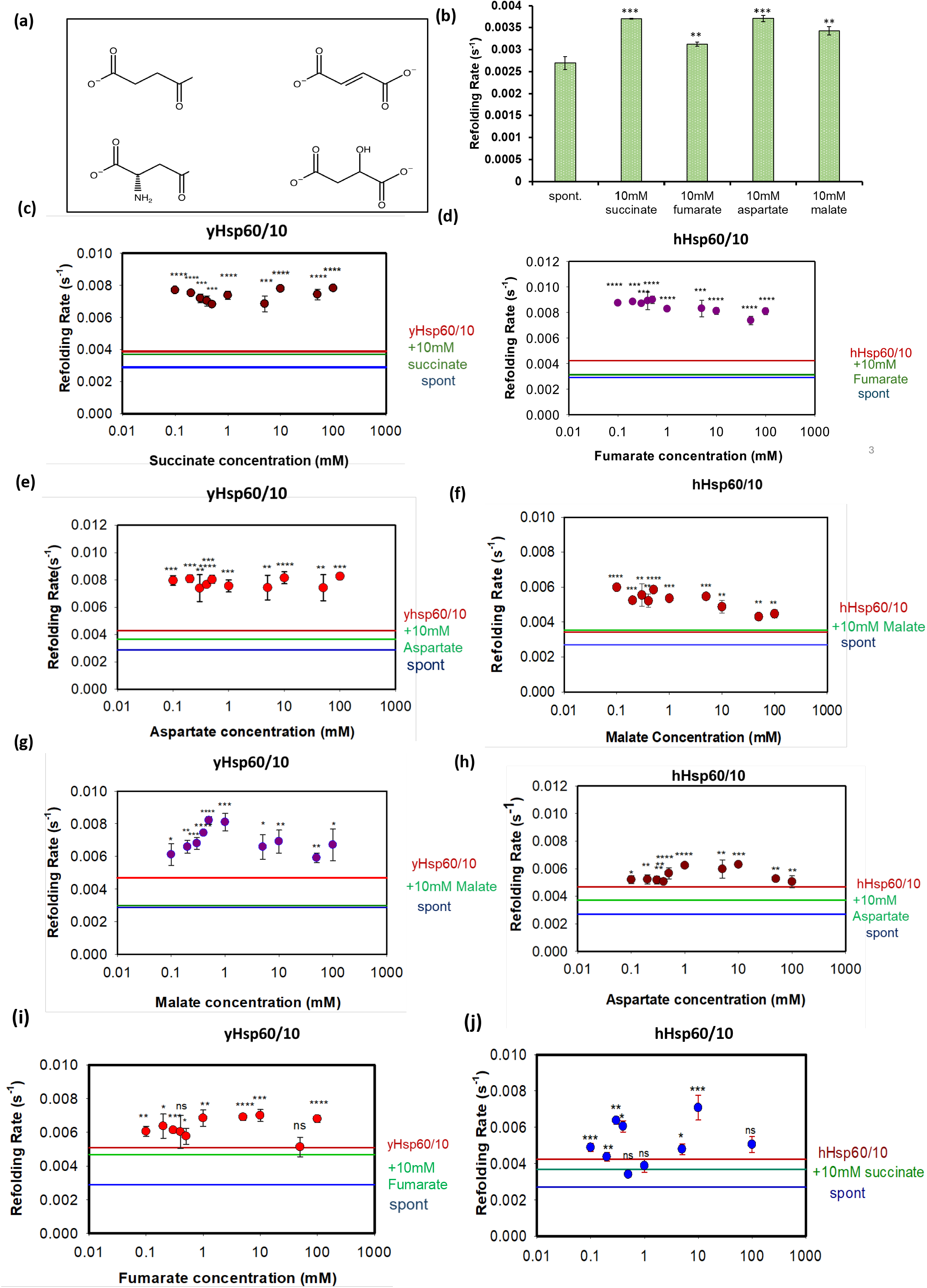
Comparison of Refolding kinetics of sGFP in the presence of small negatively charged molecules when supplemented with GroEL/ES Homologs. **(a)** Structure of succinate, fumarate, aspartate, and malate in their anioinic form. **(b)** Refolding kinetics of sGFP in the presence of 10mM small negatively charged molecules, the refolding rate is significantly not increased by small molecules compared to supplementation with GroEL/ES homologs. Rates of yHsp60/10 assisted refolding of sGFP in the presence of different concentrations of (c) succinate, (e) aspartate, **(g)** malate **(i)** fumarate. Rates of hHsp60/10 assisted refolding of sGFP in the presence of different concentrations of (d) fumarate, (h) aspartate, **(f)** malate **(j)** succinate. The blue line represents the rate of spontaneous sGFP refolding; the green line represents the rate of sGFP refolding in the presence of 10mM of small molecule, and the red line depicts the rate of sGFP refolding in the presence of yHsp60/10 or hHsp60/10.

This property could be either because of the effect of added small molecules on the folding landscape or their impact on the conformational alterations in the chaperonin itself. We checked the latter by measuring the impact of the dicarboxylic acids on the ATP hydrolysis rate of the Hsp60 or the Hsp60/10 complexes. These rates provide a sensitive indicator for alterations in the overall cycle governed by ATP binding, hydrolysis, and the allosteric changes in conformation associated with these events. The ATP hydrolysis rates did not change due to dicarboxylic acids (Figure S1A). This indicates that the small molecules likely affect the folding of the substrate than the chaperonins themselves.

To check if this is exclusive to hHsp60/10 or yHsp60/10 system, we checked the rate of GroEL/ES assisted refolding of sGFP in the presence of these small molecules; none of the small molecules increased the refolding rate further (Figure S1B). Thus, these dicarboxylic acids can synergize with hHsp60/10 or yHsp60/10 systems but not with GroEL/ES in assisting the refolding rate of sGFP. These observations posed two questions regarding the specificity of this activity: First, would these small molecules be able to collaborate with all chaperones that lack the negatively charged cavity? Second, is the synergy a specific property of negatively charged small molecules, or is it a general property of either positively charged or zwitterionic molecules?

To answer the first question, we checked if the prokaryotic homolog of GroEL/ES isolated from *Candidatus Karelsulcia muelleri* (smhEL/ES), one that does not harbour a negatively charged inner cavity, can work with the small molecules synergistically. None of the small molecules tested could increase the smhEL/ES-assisted refolding rate of sGFP (Figure 3A, B). This hinted that the synergism may be a unique property of yHsp60/10 and hHSp60/10 rather than a passive property of a less negatively charged cavity. This is further supported by the specific differences between the ability of different dicarboxylic acids to aid yHsp60/10 and hHsp60/10 assisted refolding of sGFP (Figure 2C-J).

**Figure 3.**
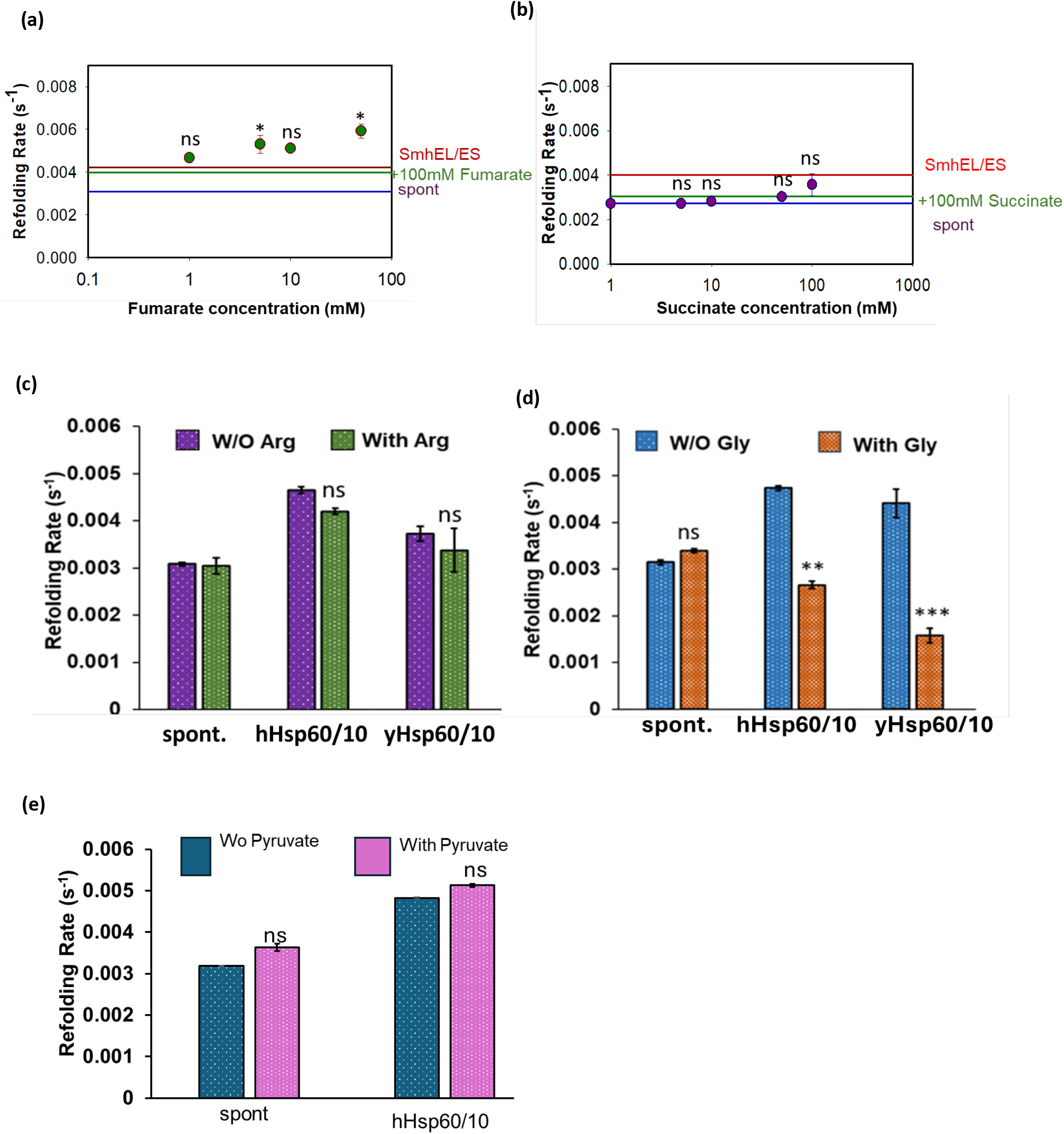
Candidatus Karelsulcia muelleri (smhEL/ES) does not cooperate with small molecules. Refolding kinetics of sGFP in the presence of small negatively charged molecules in the presence of chaperonin smhEL/ES **(a)** Fumarate at the concentration 1mM to 100mM **(b)** succinate at the concentration 1mM to 100mM; the Blue line represents the rate of spontaneous sGFP refolding, the green line represents the rate of sGFP refolding in the presence of 10mM of small molecule and the red line depicts the rate of sGFP refolding in the presence of smhEL/ES **(c)** Refolding rate of sGFP in the presence and absence of positive charged small molecule arginine **(d)** Refolding rate of sGFP in the presence and absence of glycine at the concentration of 100mM supplemented with human and yeast Hsp60/10. **(e)** Refolding rate of sGFP in the presence and absence of pyruvate at the concentration of 100mM.

To address the second question, we monitored the synergism between the chaperonins (yHsp60/10 and hHsp60/10) and either Arginine (a positively charged amino acid known to act as a chemical chaperone) or Glycine (a zwitterionic molecule also known to act as a chemical chaperone). Neither of these amino acids could increase the chaperonin-assisted refolding rate of sGFP (Figure 3C, D), indicating that this is not a general property of any charged small molecule. Additionally, a singly negatively charged molecule, pyruvate, which resembles a dicarboxylic acid structure, was unable to act like the dicarboxylic acids (Figure 3E), indicating the importance of the dual negative charge in synergizing with the eukaryotic chaperonins.

### Dicarboxylic acids cooperate with eukaryotic chaperonins to make the folding landscape like GroEL/ES assisted folding

While unidimensional comparison of spontaneous and chaperonin-assisted apparent refolding rates of substrate proteins like sGFP provide clues to the features of chaperonins required for efficient rate acceleration, it falls short of providing a more nuanced picture of the folding landscape and how the chaperonins alter it. Temperature dependence of the refolding rates allows a modified Eyring analysis of the refolding system to provide further insights into the refolding landscape (Figure S2 A, B) [22]. Given the prior knowledge that sGFP has a prominent entropic trap in its refolding landscape that GroEL/ES can remove, we asked if the synergism between the dicarboxylic acids and the eukaryotic Hsp60/10s can work as efficiently as GroEL/ES.

First, we checked if the dicarboxylic acids change the folding landscape. Although these molecules do not overtly change the refolding rate, they may change the landscape subtly, which may not be apparent in the refolding rates due to entropy-enthalpy compensation. The small molecules did not alter the entropic barrier significantly (Figure 4A). Similar observations are made for the chaperonins alone (Figure 4B). Importantly, fumarate was able to remove the entropic barrier efficiently in collaboration with hHsp60/10 (Figure 4C) and to some extent with yHsp60/10 (Figure 4D) (Table-1). Although the other combinations could accelerate the refolding rate, they could not remove the entropic trap efficiently. When acting along with yHsp60/10, most of the dicarboxylic acids decreased the entropic trap marginally while not affecting the enthalpic trap, which led to an overall decrease in the free energy of activation. Thus, the small molecules can collaborate with chaperonins to increase the folding rate and change the folding landscape of the substrate sGFP towards the same direction as GroEL/ES-assisted refolding.

**Table 1:** Summary of the kinetic parameters obtained by fitting the Eyring equation to temperature-dependent refolding data.

| Sample | $\Delta H^\#$ | $\Delta H^\# \text{ error}$ | $\rho^\#$ | $\rho^\# \text{ error}$ | $\Delta C_p^\#$ | $\Delta C_p^\# \text{ error}$ |
| --- | --- | --- | --- | --- | --- | --- |
| **Spontaneous refolding.**<br>(Sadat, A.,2022) | 4.6467 | 1.1443 | 0.0252 | 0.0483 | -0.5071 | 0.3627 |
| **GroEL/ES** | 10.3126 | 0.8199 | 873.0767 | 1197.6201 | 0.0965 | 0.2342 |
| **hHsp60/10**<br>(Sadat, A.,2022) | 0.37 | 1.05 | 1.44E-05 | 2.53E-05 | 0.42 | 0.34 |
| **10mM succinate** | 4.1907 | 0.3027 | 0.0138 | 0.007 | 0.5165 | 0.1035 |
| **10mM Fumarate** | 2.2561 | 0.5253 | 0.0004 | 0.0004 | -0.2791 | 0.1732 |
| **10mM Aspartate** | 2.7832 | 0.798 | 0.0014 | 0.0019 | -0.0779 | 0.2654 |
| **10mM Malate** | 2.7551 | 1.916 | 0.0012 | 0.0039 | -0.0513 | 0.6384 |
| **hHsp60/10+10mM succinate** | 7.1718 | 0.3463 | 3.8576 | 2.234 | 0.8165 | 0.1143 |
| **hHsp60/10+10mM fumarate** | 8.65 | 0.44 | 62.4099 | 45.9361 | -0.4181 | 0.13 |
| **hHsp60/10+10mM aspartate** | 3.7239 | 0.6628 | 0.0099 | 0.011 | -0.3883 | 0.2142 |
| **hHsp60/10+10mM malate** | 4.9088 | 0.6324 | 0.073 | 0.0773 | 0.0585 | 0.2063 |
| **yHsp60/10**<br>(Sadat, A.,2022) | 3.66 | 0.54 | 5.14E-03 | 4.68E-03 | -0.01 | 0.18 |
| **yHsp60/10+10mM succinate** | 4.7521 | 0.3698 | 0.0737 | 0.0456 | -0.117 | 0.1196 |
| **yHsp60/10+10mM fumarate** | 6.43 | 0.1 | 1.14 | 0.18 | -0.1742 | 0.0301 |
| **yHsp60/10+10mM aspartate** | 5.6593 | 2.1287 | 0.311 | 1.1083 | -0.1188 | 0.6768 |
| **yHsp60/10+10mM malate** | 5.7296 | 2.2106 | 0.3526 | 1.305 | -0.393 | 0.691 |

The parameters reported are: $\Delta H^\#$ enthalpy of activation at 298K, $\rho^\#$ is the value of activation parameter $\rho$ at 298K, and $\Delta C_p^\#$ is the change in heat capacity at constant pressure between the reactant and the transition state. Error values are standard errors obtained from fitting. All the experiments were carried out three times at each temperatures. The rates were averaged and then fitted to the modified Eyring equation. Some values used for comparison are taken from published literature from our group and are indicated by the reference. The cells are marked blue to distinguish these parameters.

**Figure 4.**
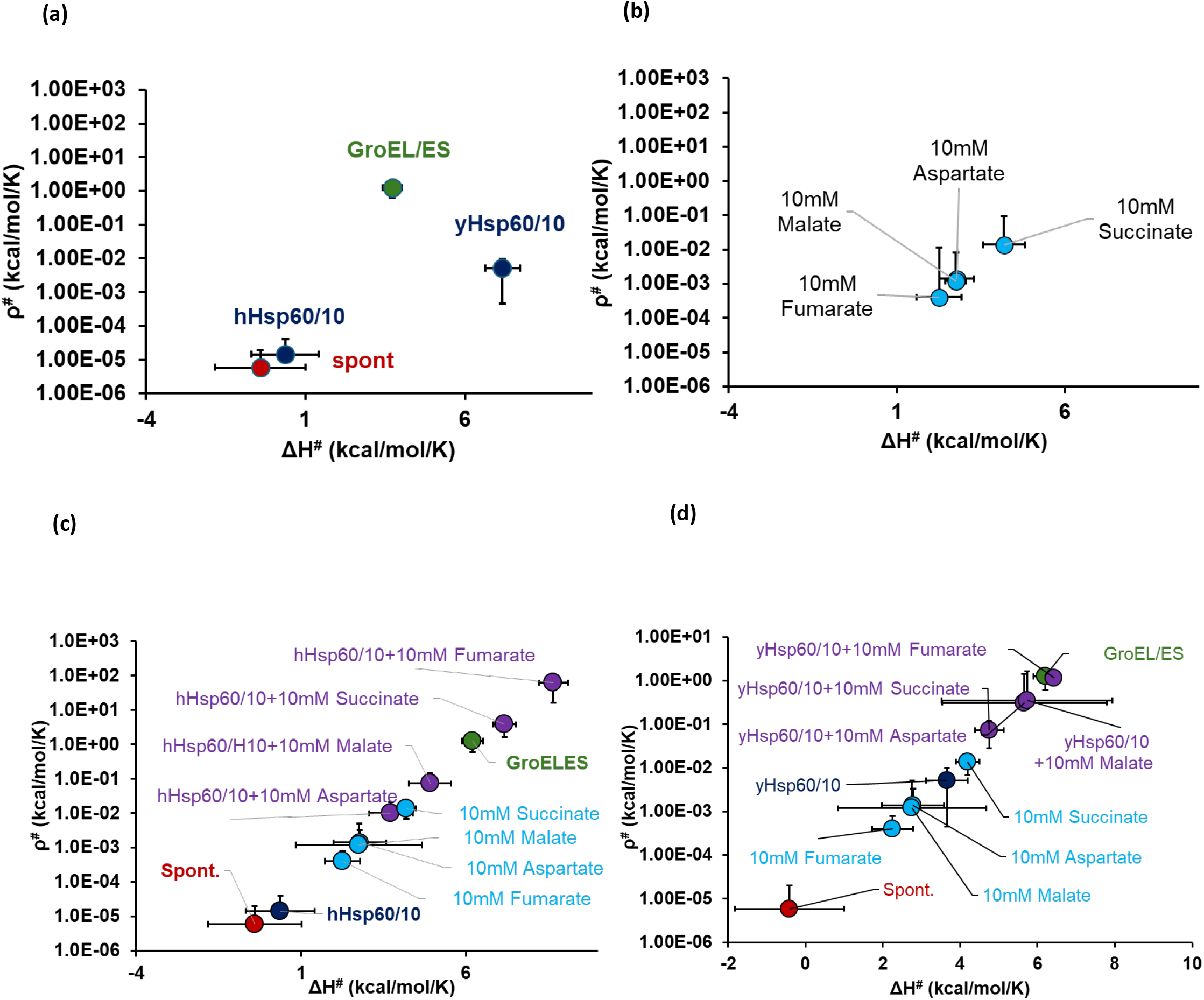
Dicarboxylic acids collaborate with eukaryotic chaperonins to mimic the folding landscape observed in GroEL/ES-assisted folding. **(a)** Thermodynamic parameters ΔH^#^ and ρ^#^ calculated in the presence of chaperonin, red closed circle-spontaneous refolding of sGFP, violet closed circle-refolding rate of sGFP in presence of yHsp60/10 and hHsp60/10, green closed circle-refolding rate of sGFP in presence of GroEL/ES **(b)** refolding rate of sGFP in presence of small molecules like succinate, fumarate, malate, aspartate, blue closed circle-**(c)** hHsp60/10 and **(d)** yHsp60/10 in presence small molecules succinate, fumarate, aspartate and malate supplementation at the concentration of 10mM in presence of chaperonin

We plotted ρ^#^ vs the folding rate (Figure 5A, B) to obtain the correlation between the ability of the chaperonins to accelerate refolding and remove the entropic trap from the folding landscape of sGFP. There is a clear correlation seen between ρ^#^ and k_f_. An increase in ρ# is associated with an accelerated refolding rate. While the kinetic analysis showed the changes in the folding landscape, the synergism between the small molecules and chaperonins is not obvious from these. To check for synergism, we calculated the expected change of refolding rate (Figure 5C) or entropic trap (Figure 5D) in the presence of small molecules and chaperonins, assuming independent action of the two. Plotting these values with the observed changes revealed that 1) the increase in folding rate showed cooperative action between chaperonin and small molecules i.e. the refolding rate was more than the refolding rate expected if the two components did not interact. 2) The entropic trap showed significant synergism between hHsp60/10 and fumarate whereas there was only marginal or no synergy for the other cases. Thus, hHsp60/10 and fumarate pair have evolved a special cooperativity that allows them to work together in removing entropic traps from folding landscapes.

**Figure 5.**
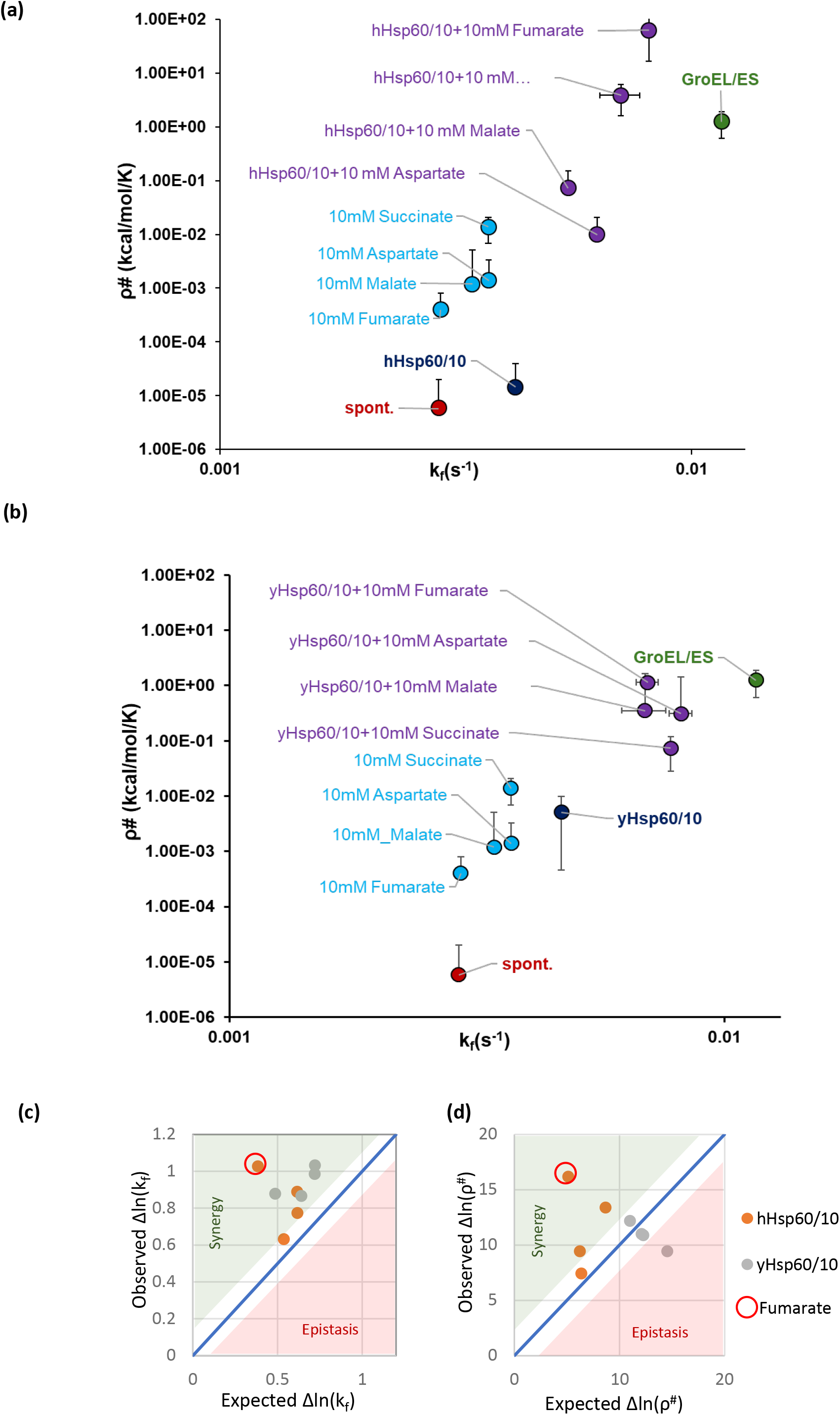
Summary of entropic parameters and the cooperativity on refolding and folding landscape. **(a)** sGFP refolding rate k_f_ and Thermodynamic parameters ρ^#^ calculated in the presence of chaperonin hHsp60/10 and small molecules. Red closed circle-spontaneous refolding of sGFP, violet closed circle-refolding rate of sGFP in presence of yHsp60/10 and hHsp60/10, green closed circle-refolding rate of sGFP in presence of GroEL/ES, blue closed circle-hHsp60/10 and yHsp60/10 in presence small molecules succinate, fumarate, aspartate, and malate supplementation at the concentration of 10mM in presence of chaperonin **(b)** sGFP refolding rate kf and Thermodynamic parameters ρ^#^ calculated in the presence of chaperonin hHsp60/10 and small molecules. **(c)** and **(d)** The expected changes in Δln(k_f_)(C) or Δln(ρ^#^)(D) is plotted vs their experimentally observed changes. The predicted change is calculated from a simple model of multiplicative changes (additive in the log scale). The diagonal line provides a visual guide to the region of no interaction between chaperonins and small molecules (i.e. observed = expected). The green triangle marks the region of synergism between chaperonin and the small molecules. In contrast, the red triangular region signifies mechanistic epistasis and, hence, similarity in the assistance mechanism between the two. The two chaperonins are marked in different colors, and Fumarate is marked separately as a particular case.

Notably, for yHsp60/10 the cooperative effect in the refolding rates (Figure 5C) but not in their action on entropic trap (Figure 5D) indicates that the cooperative effect could arise due to modulations in the enthalpic component of the folding landscape. Further investigations would reveal the exact nature of this cooperation and how this evolved in different homologs of Hsp60/10 system.

Taken together, sGFP is a substrate of GroEL/ES that benefits from the negatively charged inner cavity of the chaperonin. The eukaryotic chaperonins that have lost the inner negative charge can synergize with negatively charged small molecules, like the dicarboxylic acids, to accelerate the folding of sGFP in vitro, partially mimicking the mechanism of GroEL/ES. However, a prokaryotic homolog of GroEL/ES that lacks negative charge is not assisted by the dicarboxylic acids in folding sGFP.

## Conclusion

In this work, we find an unexpected cooperation between small molecules and a large protein machinery involved in protein folding – the class I chaperonins from eukaryotic mitochondria. While the class I chaperonins are conserved in their overall architecture through the different kingdoms of life, they show substantial differences in their central cavity [18], which is crucial to impart chaperoning environment to the encapsulated unfolded protein substrate [15] [16]. Here, we show that the cellular milieu of the chaperonin could shape the evolution of the cavity environment; some of the small molecules that are available in eukaryotic mitochondria (the dicarboxylic acids present as TCA cycle intermediates) could synergize with the mitochondrial Hsp60/10s while not cooperating with many of their prokaryotic homologs.

This cooperation could also have important implications in regulating the proteostasis capacity of the mitochondrial matrix. Changes in the concentration of dicarboxylic acids or other small molecules that synergize with the Hsp60/10s could change the activity of the chaperonin. This cooperation can potentially increase or decrease the folding capacity of the mitochondrial matrix through the regulation of metabolism that produces or consumes these metabolites. The true implication of this cooperation will require further experimentation, and we are hopeful that this work will be the basis for future investigations into the cooperation between metabolism and chaperone machinery.

## Materials and methods

### Strains

cloning experiments were employed by using E. coli DH5α strains. For the expression of arabinose-inducible pBAD GFP, we utilized BL21-DE3 Rosetta strain. BW25113 (referred as Wtk hereafter) was used to express and purify different chaperonins.

### E.coli GroEL/ES, hHsp60/10, yHsp60/10, SmhEL/ES purification

GroEL/ES and its homologs were expressed and purified using a pOFX vector containing chaperonin inserts, induced using 0.5mM IPTG at 37°C overnight. These were purified using the previous protocols [14] [23] [24].

### In-vitro refolding of spontaneous and GroEL/ES and its homologs (hHsp60/10, yHsp60/10)

Substrates were unfolded in 6M GuHCl containing buffer for 1 hour at 25°C. For refolding, a low salt buffer containing 25 mM Tris-HCl (pH 7.4), 20 mM KCl, 10 mM MgCl_2_, and 2 mM DTT (pH 7.4) was used. Substrate refolding was initiated by 100-fold dilution in buffer at 25°C to the final substrate concentration of 200nM. Chaperone-assisted refolding was done in the presence of 400nM GroEL (tetradecamer) and 800nM GroES (heptamer), such that the ratio of substrate:GroEL:GroES::1:2:4. The refolding is assisted by adding 200mM ATP. The GFP fluorescence readout was taken in Fluorolog-3 (Horiba) spectrofluorometer with excitation at 480nm with a slit width of 2nm and emission at 515nm with a slit width of 5nm [25].

### In-vitro refolding of spontaneous and GroEL/ES and its homologs (hHsp60/10, yHsp60/10) with supplementation of small molecules

For *in-vitro* refolding with supplementation of small negatively charged molecules, succinate, fumarate, aspartic acid, malic acid pyruvate, arginine, and glycine were used. As mentioned above, the refolding assay was done at different concentrations of 0.1-100mM of small molecules added in the refolding reaction.

### ATPase assay

Rate of ATP hydrolysis was calculated using an enzyme coupled ATP consumption assay [26]. 2mM phosphoenol pyruvate, 20/30 units/mL pyruvate dehydrogenase/ lactate dehydrogenase enzyme, 1mM NADH, and 1mM ATP were added in a reaction mixture in low salt buffer and incubated at 25°C for 5 minutes to remove any ADP present. 10mM fumarate was added where mentioned. 2μM of GroES or Hsp10 was added to the reaction mixture, and the reaction was started by addition of 1μM of GroEL or Hsp60. Decrease in absorbance was measured at 340nM. The rate of decrease in NADH is equal to the rate of ATP hydrolysis and plotted as ATPase rate (μM/min per μM of GroEL) [18].

### Temperature-dependent refolding analysis by using the Arrhenius equation

To characterize the thermodynamic parameters that govern the barrier between the refolding intermediate I_1_ and the transition state (TS) of folding, we employed the following equation, which is fundamentally outlined in [22].

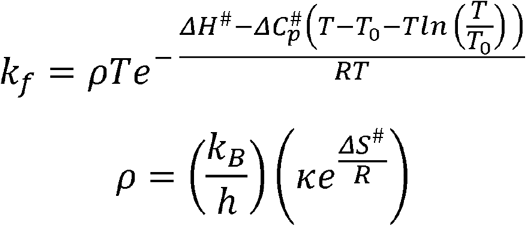

k_f_ is the refolding rate, κ is the transmission factor, indicating the proportion of activations that lead to the formation of the native state (N), k_B_ is the Boltzmann constant, T is the temperature of the refolding reaction, *h* is the Planck constant, ΔH^#^, ΔS^#^ and ΔC_p_^#^ are the differences in enthalpy, entropy and heat capacity between the transition state (TS) and intermediate I_1_. R is the universal gas constant, T is the temperature of the refolding reaction, and T^0^ is the reference temperature at which the parameters are determined (298.15 K).

The equations underwent standard nonlinear regression fitting using R or Octave. Fitting involved adjusting the starting parameters by a factor of four within the expected range reported for globular proteins [22]. Fitting was considered satisfactory when the r-squared values exceeded 0.9 and the dependencies for the various floated parameters during fitting were above 0.9. The floated parameters during fitting included ρ, ΔCp^#^, and ΔH^#^.

### Flow cytometry

pBAD-sGFP-mCherry plasmid was transformed in the background of Wtk cells with a pOFX vector containing GroEL/ES, hHsp60/10, and yHsp60/10. Secondary culture was set using overnight grown cultures. At OD ∼0.6, overexpression of chaperonins was induced using 0.5mM IPTG for 45 min, and then overexpression of sGFP was induced using 0.1% arabinose for 3 hours at 37°C. Cells were diluted in 1X PBS post-induction and kept at 37°C for 30 min for mCherry maturation. Flow cytometry data was collected in BD LSR-II and analysed in MATLAB.

### Quantification and statistical analysis

Statistical analysis for quantification utilized the student’s t-test and an R package for non-linear regression. MATLAB was employed for the analysis of flow cytometry data.

## Acknowledgment

The work in the KC lab was supported by a Swarna Jayanthi fellowship Grant from DST(DST/SJF/LSA-01/2015-16). Instrument support was also obtained from CSIR and Wellcome Trust – DBT India Alliance. DG and MK acknowledge UGC for their fellowship. We also acknowledge Aseem Chaphalkar for helping in refolding assays.

## Author contribution

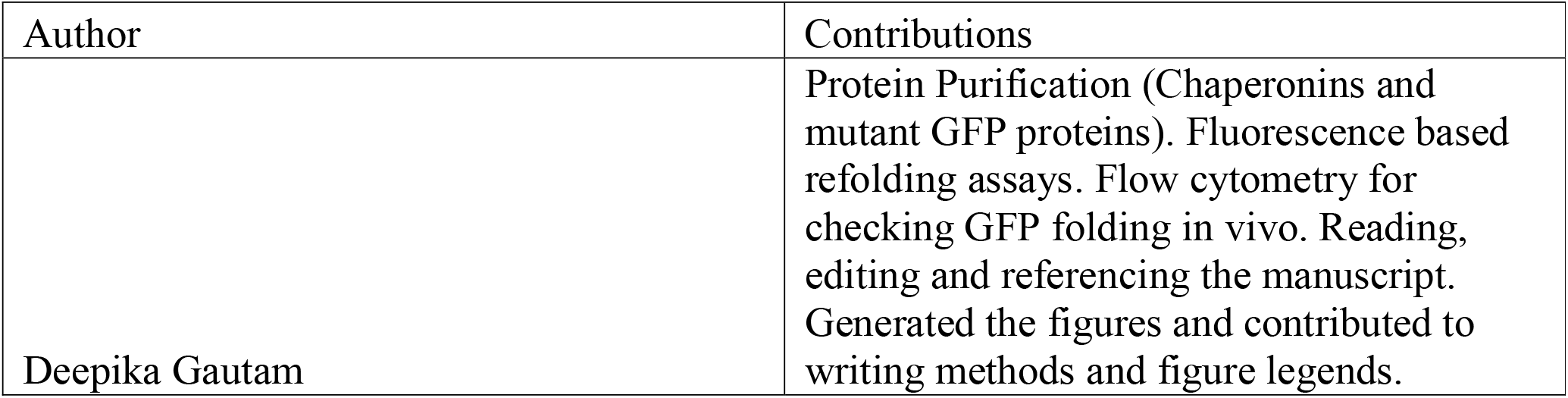
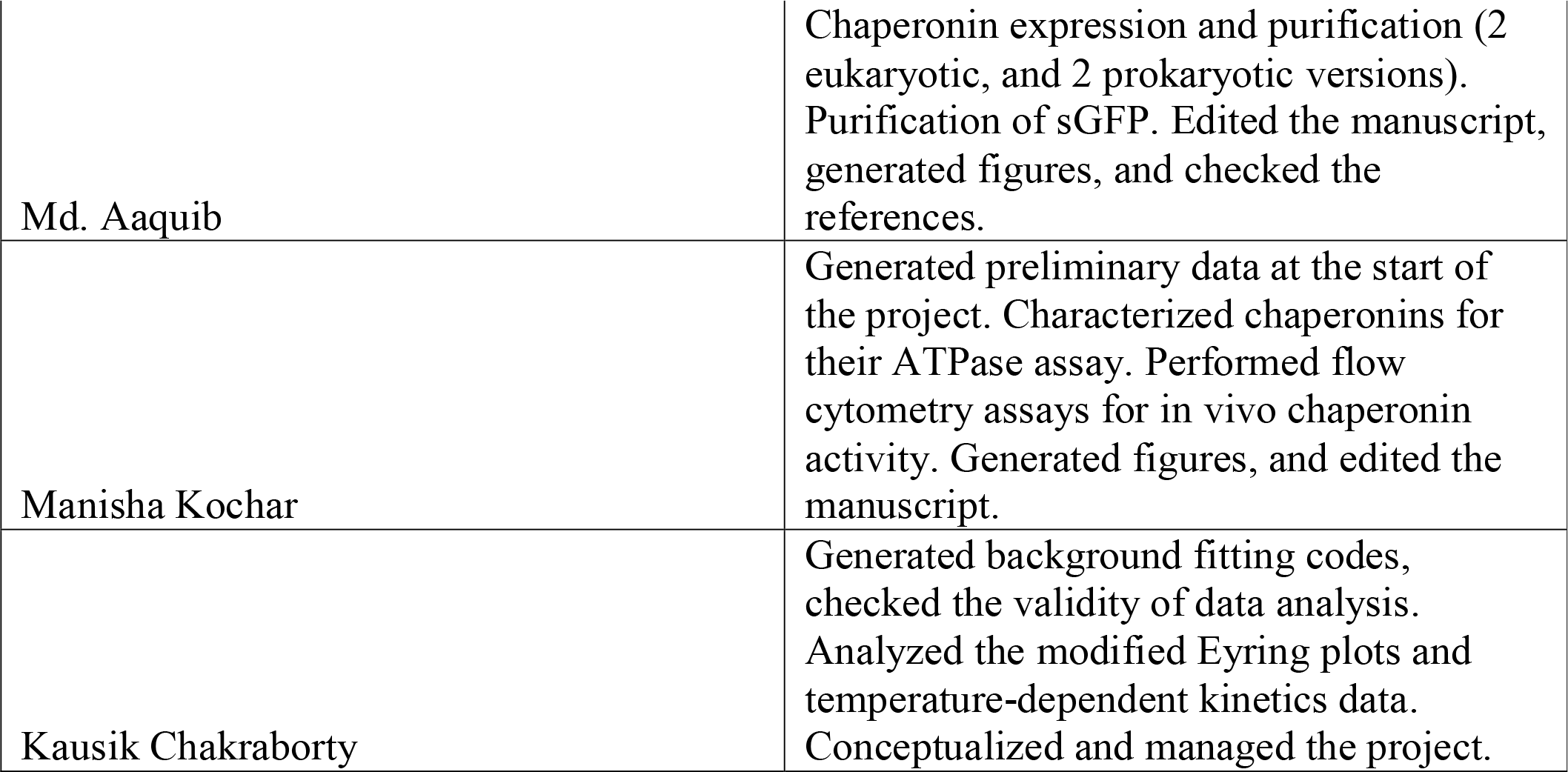

## Conflict of interest

The authors declare no competing interests.

## Notes

### Competing Interest Statement

The authors have declared no competing interest.

