## Supplemental Information for "Dicarboxylic acids synergize with yeast and human Hsp60/10 systems to mimic GroEL/ES"

**Figure Legends**

**Figure S1**
**a)** Rate of ATP hydrolysis in the presence or absence of 10 mM fumarate with GroEL (Hsp60) alone or GroEL/ES (Hsp60/10).

**b)** Refolding rate of sGFP in the presence of GroEL/ES, and supplemented with small molecules succinate and fumarate at the concentration of 10mM.

**Figure S2**

**a)** Temperature-dependent refolding kinetics of sGFP with hHsp60/10 supplemented with **(a)** 10mM succinate, **(b)** 10mM fumarate **(c)** 10mM aspartate, **(d)** 10mM malate. Red line depicts 95% of the prediction band, the blue line shows 95% of the confidence band, and the black line depicts the best fit of refolding rate in different temperatures. The x-axis shows different temperatures from 15°C to 35°C, and the y-axis shows the refolding rate of substrate protein sGFP.

**b)** Temperature-dependent refolding kinetics of sGFP with yHsp60/10 supplemented with (**a)** 10mM succinate, (**b)** 10mM fumarate (**c)** 10mM aspartate, (**d)** 10mM malate. Red line depicts 95% of the prediction band, the blue line shows 95% of the confidence band, and the black line depicts the best fit of refolding rate in different temperatures. The x-axis shows different temperatures from 15°C to 35°C, and the y-axis shows the refolding rate of substrate protein sGFP.

**Figure S1**

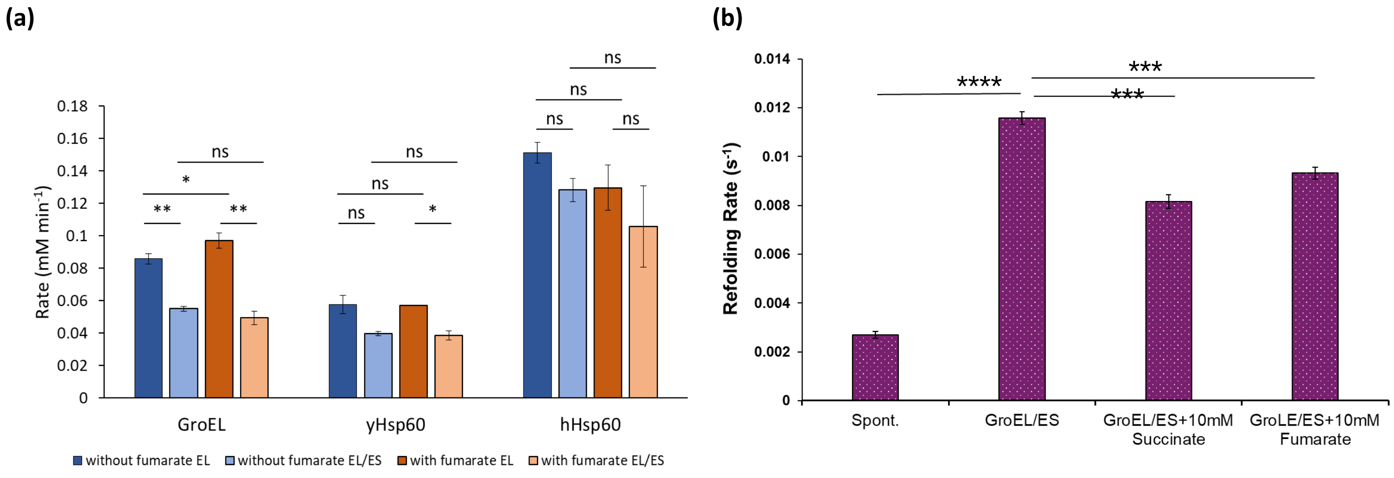
(a)
(b)
0.18
0.16
0.14
0.12
0.1
0.08
0.06
0.04
0.02
0
Rate (mM min-1)
ns
*
**
**
ns
ns
ns
*
ns
ns
ns
ns
GroEL
yHsp60
hHsp60
without fumarate EL
without fumarate EL/ES
with fumarate EL
with fumarate EL/ES
0.014
0.012
0.01
0.008
0.006
0.004
0.002
0
Refolding Rate (s-1)
****
***
***
Spont.
GroEL/ES
GroEL/ES+10mM Succinate
GroLE/ES+10mM Fumarate

**Figure S2**

**(a)**

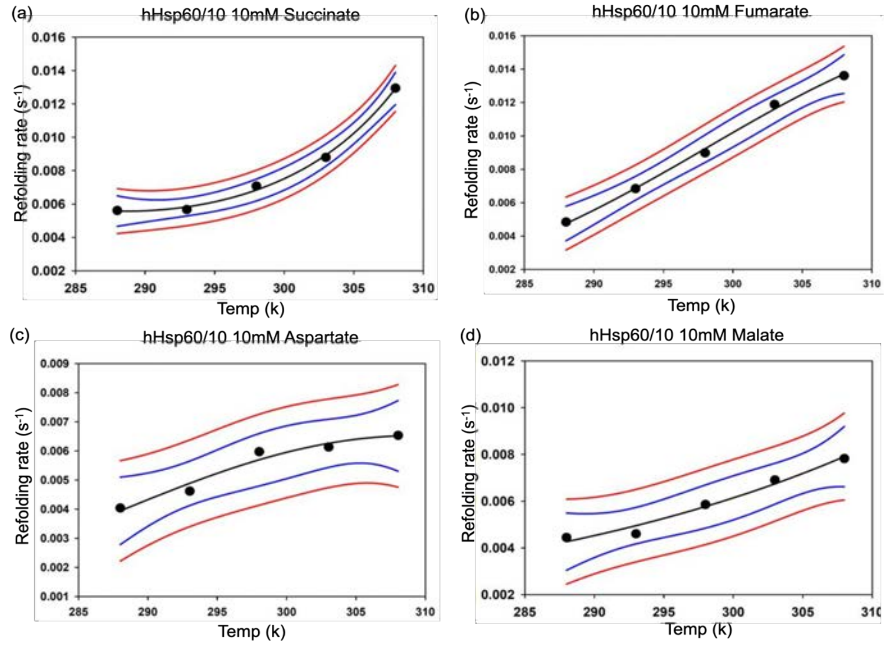
(a)
hHsp60/10 10mM Succinate
(b)
hHsp60/10 10mM Fumarate
(c)
hHsp60/10 10mM Aspartate
(d)
hHsp60/10 10mM Malate
Refolding rate (s-1)
Temp (k)

**(b)**

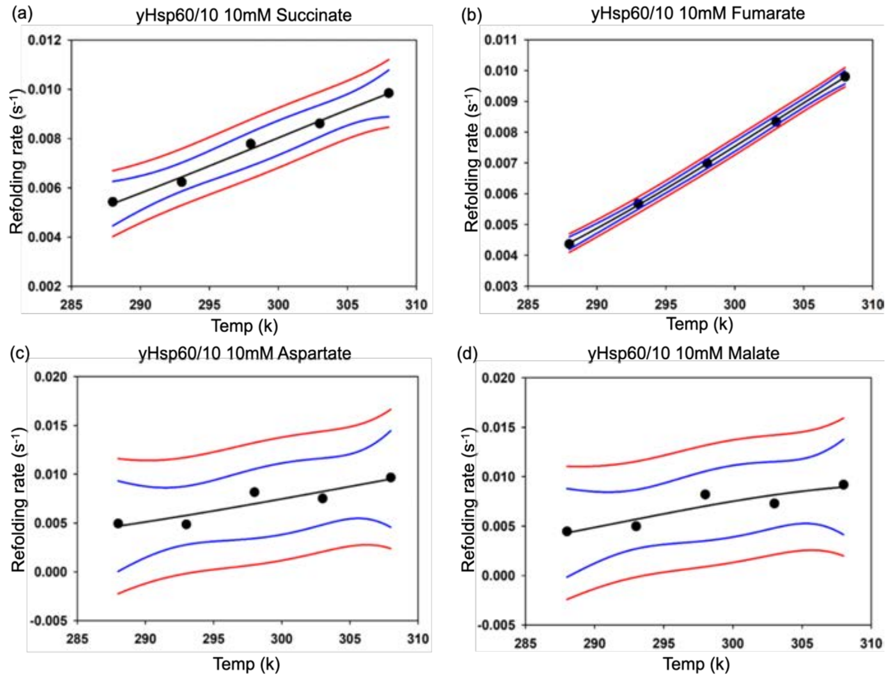
(a)
yHsp60/10 10mM Succinate
(b)
yHsp60/10 10mM Fumarate
(c)
yHsp60/10 10mM Aspartate
(d)
yHsp60/10 10mM Malate
Refolding rate (s-1)
Temp (k)
